# Bei Mu Gua Lou San facilitates mucus expectoration by increasing surface area and hydration levels of airway mucus in an air-liquid-interface cell culture model of the respiratory epithelium

**DOI:** 10.1101/2023.01.31.526405

**Authors:** Silvia Groiss, Ina Somvilla, Christine Daxböck, Manuela Stückler, Elisabeth Pritz, Dagmar Brislinger

**Author notes:** Correspondence to*: PD Dr. Dagmar Brislinger, MSc, Division of Cell Biology, Histology and Embryology, Gottfried Schatz Research Center, Medical University of Graz, Neue Stiftingtalstraße 6/II, 8010 Graz, Austria, *E-Mail address.

## Abstract

Bei Mu Gua Lou San (BMGLS) is an ancient formulation known for its moisturizing and expectorant properties, but the underlying mechanisms remain unknown. We investigated dose-dependent effects of BMGLS on its rehydrating and mucus-modulating properties using an air-liquid-interface (ALI) cell culture model of the Calu-3 human bronchial epithelial cell line and primary normal human bronchial epithelial cells (NHBE), and specifically focused on quantity and composition of the two major mucosal proteins MUC5AC and MUC5B.

ALI cultures were treated with BMGLS at different concentrations over three weeks and evaluated by means of histology, immunostaining and electron microscopy. MUC5AC and MUC5B mRNA levels were assessed and quantified on protein level using an automated image-based approach. Additionally, expression levels of the major mucus-stimulating enzyme 15-lipoxygenase (ALOX15) were evaluated. BMGLS induced dose-dependent morphological changes in NHBE but not Calu-3 ALI cultures that resulted in increased surface area via the formation of herein termed intra-epithelial structures (IES). While cellular rates of proliferation, apoptosis or degeneration remained unaffected, BMGLS caused swelling of mucosal granules, increased the area of secreted mucus, decreased muco-glycoprotein density, and dispensed MUC5AC. Additionally, BMGLS reduced expression levels of MUC5AC, MUC5B and the mucus-stimulating enzyme 15-lipoxygenase (ALOX15).

Our studies suggest that BMGLS rehydrates airway mucus while stimulating mucus secretion by increasing surface areas and regulating goblet cell differentiation through modulating major mucus-stimulating pathways.

## Introduction

Chronic respiratory diseases (CRDs) are a class of widespread multifaceted airway disorders marked primarily by increased airflow obstruction due to hyperproduction of airway mucus and inflammation with a global prevalence of over 1 billion cases; tendencies increasing (Bousquet et al., 2010; Holgate, 2012; Rabe & Watz, 2017). Bronchodilators and corticosteroids are the current backbone of CRD treatment but show considerable side effects (Matera et al., 2011; Wenzel, 2012), while antibody therapies are limited in patient responsiveness (Fahy et al., 1997; Hanania et al., 2011). The need for new treatment perspectives redirects research towards traditional herbal medication that is widely believed to offer novel and effective possibilities for patients with CRDs.

In China, the ancient formulation Bei Mu Gua Lou San (BMGLS) has been administered over 2000 years to treat CRDs through moisturizing airway mucus and mitigating inflammation (Chen et al., 2020; Köfers, 2009). Bulbs of the genus *Fritillaria cirrhosa* D.Don are the main ingredient and reported as a potent antitussive and anti-asthmatic herbal drug in numerous studies (Lin et al., 2001; Zhou et al., 2010). Further ingredients elicited antioxidant, antibiotic, anti-inflammatory, neuroprotective and antiviral properties (Nyakudya et al., 2014; Ozaki et al., 1996; Rios, 2011; Shaw et al., 2005; Y. Zhao et al., 2017). However, detailing the underlying molecular mechanisms of the composite formulation of BMGLS on CRDs has not been addressed so far.

CRD development is often preceded by epithelial reorganization. A functional respiratory epithelium intercalates various cell types among which ciliated, goblet and club cells constitute the main components that mediate mucociliary clearance (MCC) (Bustamante-Marin & Ostrowski, 2017).

The air-surface-liquid (ASL) that mediates MCC comprises the displaceable mucous layer that rests on the perciliary layer (PCL), which interacts with apical cilia. Aided by coordinated ciliary beating, these components propel the mucous layer including trapped particles towards the proximal trachea for expectoration. Increased levels of the mucosal proteins MUC5AC and MUC5B often hamper effective MCC in CRDs by altering the hydration state and composition of the ASL, eventually aggravating mucosal viscidity (Hamed & Fiegel, 2014; Jackson et al., 2020; Kesimer et al., 2017; Matthews et al., 1963). Detailed studies on the role of both mucins in MCC using knock-out studies in mice revealed a beneficial yet negligible role for MUC5AC, while it clearly highlighted MUC5B as imperative (Roy et al., 2014). Absence of MUC5B accumulated materials such as bacteria due to non-resolving infection causing apoptotic immune cells such as macrophages to concentrate. In fact, MUC5AC has been shown to increase in pathologic conditions while MUC5B rather decreases, which clearly emphasizes the importance of a physiologic ratio of these two mucins in balancing immune homeostasis (Rose & Voynow, 2006; Roy et al., 2014).

In addition, accumulating levels of airway mucus foster goblet cell metaplasia and increase mucus production by activating 15-lipoxygenase (ALOX15), another key player in goblet cell differentiation and mucus production (Zhao et al., 2009; J. Zhao et al., 2017). This is of particular interest since ALOX15 was inhibited by a herbal formulation showing anti-inflammatory properties similar to BMGLS (Qi et al., 2019; Zeng et al., 2011).

Therefore, the present study investigates the effects of BMGLS on goblet cell metaplasia (GCM) and goblet cell hyperplasia (GCH) on mucus production with special emphasis on the mucosal proteins MUC5AC and MUC5B in an air-liquid-interface (ALI) cell cultures system of Calu-3 and primary normal human bronchial epithelial cells (NHBE). We assessed cellular integrity and morphology including the differentiation state of key cellular components following three weeks of BMGLS application by immunostaining and electron microscopy, and evaluated the mucus quantity and composition on mRNA and protein level using our recently developed automated image-based analysis approach (Groiss et al., 2022). We further explored whether the potential changes in mucus quantity may attribute to regulation of key mucus-stimulating pathways.

Overall, we hypothesize that BMGLS fosters mucus expectoration by increasing the hydration state of airway mucus potentially by increasing the surface area for facilitated expectoration and may therefore proof beneficial as a supportive measure to conventional treatment.

## Materials and Methods

The hydrophilic concentrate of BMGLS used within this study was purchased from Conforma NV (Belgium) that used the method of Dynafyt for BMGLS production, and complies with European GMP and Chinese Pharmacopeia standards (*Pharmacopoeia of the People’s Republic of China: Pharmacopoeia of the Ministry of Health of the People’s Republic of China*., 2015). The ingredients are listed in Table 1 (www.theplantlist.org). The correct botanical identity of the components was tested using microscopy, thin-layer chromatography (TLC) and high-performance layer chromatography (HPLC) as reported in the Certificate of Analysis. Additional tests were performed to preclude contamination with harmful substances (e.g., aflatoxins, aristolochid acid, bacteria, heavy metals, pesticides, pyrrolizidine alkaloids and sulfur dioxide). The solvent solution of the hydrophilic concentrate (HCS; 30 % glycerol, 0.1 % propylene glycol, 0.1 % methyl-4-hydroxybenzoate, 0.05 % propyl-4-hydroxybenzoate), as indicated by Conforma NV, was used as solvent control in all experiments. In this study, we used concentrations of 0.07%, 0.15%, 0.3%, 0.6%, 1.25%, 2.5% and 5% (v/v) of the hydrophilic concentrate BMGLS in PneumaCult™ ExPlus or PneumaCult™ ALI medium (Stemcell Technologies, Vancouver, Canada) supplemented with MycoZAP (Lonza, Basel, Switzerland), and Anti-Anti (Gibco, Dublin, Ireland) as instructed by the supplier as well as 40 μg/ml ticarcillin (Merck, Darmstadt, Germany).

**Table 1.** Herbal formulation of BMGLS. Herbal ingredients and their proportion in BMGLS including traditional usage and effects reported in literature. Major active compounds are given.

| Herb | % of crude drug | Traditional usage | Effect acc. To Literature | Major active compounds | Reference |
| --- | --- | --- | --- | --- | --- |
| *Fritillaria cirrhosa* D.Don<br>(dried bulb of *Fritillaria cirrhosa*) | 23 | Clears heat, moistens the lung, transforms mucus, stops cough | antitussive, expectorant, anti-inflammatory, anticancer, anti-oxidative | alkaloids, saponins, imperialine, nucleosides, terpenoids, sterols, ... | (Chen et al., 2020; Köfers, 2009; Lin et al., 2001; Wang et al., 2016; Wang et al., 2011; Zhou et al., 2010) |
| *Trichosanthes kirilowii* Maxim.<br>(dried fruit and root of *Trichosanthes kirilowii*) | 15.4 each | Clears heat, Moistens dryness, regulates Qi, clears mucus | expectorant, anti-inflammatory, anti-oxidant, ribosome-inactivating, Ca<sup>2+</sup> scavenging | trichosanthin, Karasurin, Agglutinin, beta-sitosterol, ... | (Ozaki et al., 1996; Shaw et al., 2005; Yu et al., 2018) |
| *Poria cocos* Wolf<br>(inner part of the dried sclerotia of *Poria cocos*) | 15.4 | Strengthens the spleen, guides fluids | tonic, anti-inflammatory, diuretic, ameliorating phlegm and edema, anti-tumor | pachymic acid, tumulosic acid, eburicoic acid, ... | (Nie et al., 2020; Rios, 2011) |
| *Citrus reticulata* Blanco<br>(dried exocarp of *Citrus reticulata*) | 15.4 | Protects stomach, eliminates dampness And phlegm, transforms mucus | antioxidant, anti-tumor | hesperidin, nobelitin, tangeretin, ... | (Y. Zhao et al., 2017) |
| *Platycodon grandiflorus* (Jacq.) A.DC.<br>(dried root of *Platycodon grandiflorus*) | 15.4 | Promotes lung-Qi-flow | cytotoxic for tumor cells, antiviral, neuroprotective, antioxidant, immune-modulative | saponins, flavonoids, phenolic acids, ... | (Ji et al., 2020; Nyakudya et al., 2014) |

### Calu-3 cells and culture

The Calu-3 human bronchial epithelial cell line was provided from the Center for Medical Research (Medical University of Graz, Austria, MUG) and used between passages 20 and 40. Cells were cultured in Minimum Essential Medium (Gibco; MEM) completed with 10% fetal bovine serum (HyClone, Lot no RC35965; FBS), 100 μM penicillin-streptomycin (Sigma-Aldrich) and 2 mM L-Glutamine (Gibco), and maintained in a humidified, 5% CO_2_-95% atmospheric air incubator at 37°C. Medium was exchanged twice per week and cells were passaged weekly at a 1:3 split ratio by using a trypsin-EDTA solution.

### Specimen collection

Tracheal tissue was obtained from organ donors from the Division of Transplantation Surgery or during routinely performed autopsies from the Institute of Pathology (MUG) in accordance with the Austrian law BGBl. 1 Nr. 108/2012 following approval of the MUG ethical committee (EK30-377ex17/18). Donors diagnosed with bronchial carcinoma, acute or chronic pulmonary disease or current nicotine abuse were excluded. The mean age was 67.17 ± 18.17 years for female (19 samples) and 63.52 ± 15.30 for male (22 samples) donors.

### Isolation of NHBE cells

NHBE cells were isolated as previously described (Groiss et al., 2022). Briefly, the tracheal tissue was washed in sterile 0.9% NaCl / 3% penicillin-streptomycin before mechanically removing soft tissue or lung parenchyma and incubation in MEM (Gibco) supplemented with 1x MycoZAP (Lonza), 1x Antibiotic-Antimycotic (Anti-Anti, Gibco), 40 μg/mL ticarcillin (Merck), 500 μg/mL dithiothreitol and 10 μg/mL DNase (in PBS) for 4h at 4 °C followed by incubation in the aforementioned medium supplemented with collagenase solution (185 U/ml collagenase type II, 2 mg/ml BSA, 0.5 mM calcium chloride, 1 U amino acids 100x, 5% FCS in DMEM) and 10 μg/ml DNase (Gibco) over night at room temperature (RT). The epithelial cells were gently scraped off from the luminal surface of the trachea using fresh MEM and suspended in MEM supplemented with 5x Anti-Anti solution for incubation for 2h at 37°C to remove any residual contaminants. The cells were then suspended in PneumaCult™ ExPlus medium (Stemcell) supplemented with 1x Anti-Anti, 1x MycoZAP and 40 μg/mL ticarcillin and seeded into gelatin-coated cell culture flasks.

### ALI cultures of Calu-3 and NHBE cells

Following cell harvesting (Calu-3) or isolation (NHBE), the cells were seeded into fibronectin-coated (2 μg/cm^2^; air dry for 1 hour; Sigma-Aldrich) transwell cell culture supports (VWR, Dublin, Ireland) at a density of 3.5 × 10^5^ cells (Calu-3) or 3×10^5^ cells (NHBE) per 12-well insert/cell culture support in 0.5 mL medium with 1 mL medium added to the basolateral chamber and incubated at 37 °C and 5% CO_2_. As suggested by the manufacturer, the apical compartment was cleared from medium after three days and PneumaCult™ ALI medium, including BMGLS, was henceforth supplied only in the basal compartment with medium changes every other day.

### Epithelial height measurement

The epithelial height was measured using ImageJ and OlyVIA (Olympus) software on three different locations per membrane and sample and converted to μm according to the specifications of the microscope. The epithelial height of the ALI cultures was tracked throughout the 3 weeks’ cultivation period.

### Transepithelial electric resistance

The trans-epithelial electric resistance (TEER) was tracked using the EVOM voltohmmeter equipped with STX-2 chopstick electrodes (World Precision Instruments, Stevenage, UK) during wet (days −3, −2 and −1) and ALI culture (days 0, 3, 7, 10, 14 and 21). The apical and basolateral side contained 0.5 and 1 mL of ALI medium, respectively, during the measurement. TEER values were corrected for the blank, as posed by the fibronectin-coated non-cellularized insert, and the surface area.

### Cytotoxicity assessment of BMGLS

The hydrophilic concentrate BMGLS was tested for cytotoxicity using a 3-(4,5-dimethylthiazol-2-yl)-2,5-diphenyl tetrazolium bromide (MTT) assay (EZ4U Assay, Biomedica, Vienna, Austria) according to manufacturer’s instructions. The Calu-3 cells or primary NHBE cells were seeded in a 96-well culture plate (Nunc, ThermoFisher Scientific, San Francisco, USA) at a density of 5×10^4^ cells/well, let adhere for 48h and incubated for 24h with BMGLS at concentrations of 0.07%, 0.15%, 0.3%, 0.6%, 1.25%, 2.5%, and 5% (v/v) with HCS at 5% (v/v) as solvent control. The absorbance of dissolved formazan was measured at 584 nm (Flustar Optima, BMG Labtechnology, Offenburg, Germany) and is given as percentage of the untreated control.

### BMGLS treatment of ALI cultures

BMGLS was supplied basolateral at concentrations of 0.15%, 0.3% and 0.6% (v/v) in PneumaCult™ ALI culture medium, supplemented as described above, with HCS at 0.6% (v/v) and medium only as the controls over three weeks starting with the air-lift of the ALI culture (post 3 days of wet culture). All cultures were kept at 37°C and 5% CO_2_.

### Sample processing for histology and automated image-based analysis

ALI cultures were fixed using Carnoy’s reagent (ethanol, chloroform, glacial acetic acid; 6:3:1, v/v) for 20 min, flushed in PBS, paraffin-embedded using an Excelsior™ AS Tissue Processor (ThermoFisher Scientific), sectioned to 4 μm and mounted on SuperFrost Plus™ slides before dewaxing using Histolab Clear (Histolab^®^, Askim, Sweden) and rehydration in a descending series of ethanol (100 %, 96 %, 70 % and 50 %). Antigen retrieval was done using 0.1 M sodium citrate buffer solution (pH 6) for 15 min at 93° C by using a KOS Microwave Multifunctional Tissue Processor (Milestone, Sorisole, Italy).

Alcian blue (AlB) staining was conducted by incubation in 3% acetic acid for 3 min before incubation in 1% alcian blue (in 3% acetic acid) for 10 min and flushed in 3% acetic acid and dH_2_O. Nuclear fast red (NFR) was used for counterstaining. No NFR was used for combinational staining of AlB and IF. Here, the slides were directly subjected to background blocking using the Ultra Vision Protein Block for 5 min (ThermoFisher Scientific) and exposed to primary and secondary antibodies for 45 min and 30 min at RT, respectively. 4′,6-diamidino-2-phenylindole (DAPI) was used for nuclei counterstaining (1:2000). Samples were mounted using ProLong Gold Antifade Reagent (both ThermoFisher Scientific).

Immunostaining was performed using the UltraVision Detection System HRP Polymer Kit (ThermoFisher Scientific) following the manufacturer’s instructions. Mouse and/or rabbit IgG (1:1000, 1μg/mL, diluted in antibody diluent, Agilent Technologies) served as negative control. Antibody specifications are given in Table 2.

**Table 2.**
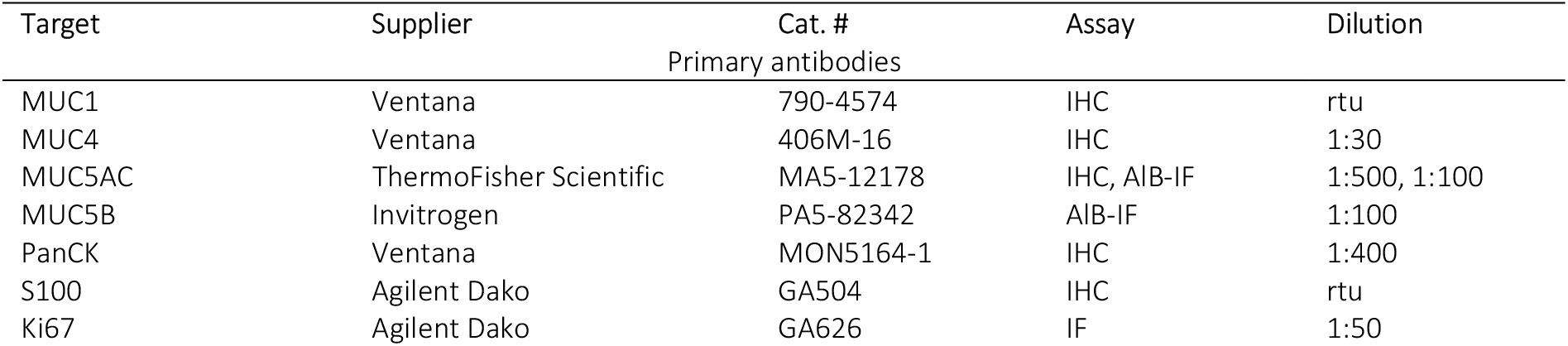
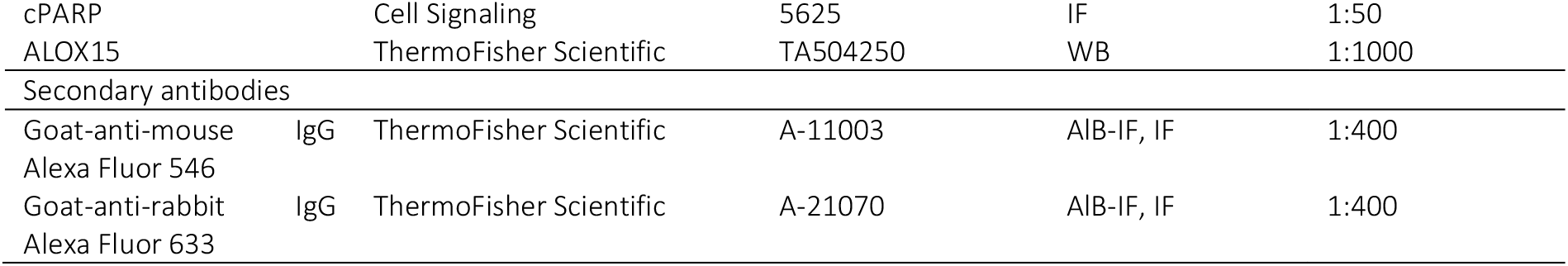
Antibody specificities for IHC and IF.

### Scanning (SEM) and transmission (TEM) electron microscopy

The ALI cell culture was fixed in 2.5% (w/v) glutardialdehyde and 2% (w/v) paraformaldehyde in 0.1 M cacodylate-buffer for 30 min covering both sides of the membrane, followed by post-fixation in 2% (w/v) osmium tetroxide in 0.1 M cacodylate-buffer for 30 min and dehydration in graded series of ethanol. SEM samples were critical point dried (CPD 030, Bal-Tec, Balzers, Liechtenstein), sputter coated with gold/ palladium (SCD 500, Bal-Tec, Balzers, Liechtenstein) and analyzed using a Zeiss Sigma 500 field emission scanning electron microscope (Zeiss, Oberkochen, Germany).TEM samples were embedded in TAAB epoxy resin (Agar Scientific Ltd., Essex, England) and ultrathin sections (70nm) were cut with a UC7 Ultramicrotome (Leica Microsystems, Vienna, Austria) stained with platin blue and lead citrate. The ultrathin sections were investigated using a Zeiss EM 902 transmission electron microscope (Zeiss).

### Image acquisition and automated mucus analysis

Histological and immunostained images were captured by bright field and fluorescence microscopy using a Leica CRT6000 (Leica Microsystems, Wetzlar, Germany) or an Olympus BX63 microscope with a DP73 camera and the DP2-TWAIN software (version 2.1, all Olympus, Tokyo, Japan, if not stated otherwise). The open source programs GNU Image Manipulation Program (GIMP, version 2.8.22, http://www.gimp.org) and Cell Profiler (version 3.1.5., (Carpenter et al., 2006)) were used for image and graphical data analysis. Cell Profiler uses a number of consecutive modules to generate an algorithm known as pipeline to detect and analyze target areas. The pipelines for analysis of airway mucus and mucosal protein content were previously described in Groiss et al., 2021 (Groiss et al., 2022). The Cell-Profiler pipeline used within this study including template data sets are available at http://cellprofiler.org. In short, the mucosal area as indicated by AlB staining in bright field images was identified and used to calculate the fluorescence signal intensity of the mucosal proteins within that AlB area. Downstream calculations to assess the area and optical density of both AlB and the mucosal proteins, respectively, were conducted using the same pipeline.

### RNA isolation and quantitative reverse transcription PCR

Total RNA was isolated using the peqGOLD Total RNA Kit (Peqlab, Erlangen, Germany), tested on concentration and integrity using an ND-1000 spectrophotometer (NanoDrop Technologies, Wilmington, USA) and reverse transcribed with the ReverseAid First Strand cDNA Synthesis Kit (ThermoFisher Scientific). mRNA expression levels of target genes were analyzed by quantitative reverse transcription PCR (RT-qPCR) using the iTaq™ Universal SYBR® Green Supermix on a CFX384™ Real-Time System (both BioRad, Hercules, CA, USA) including melting curve analysis for quality assessment. Primers were purchased from Microsynth (Vienna, Austria; MUC5AC forward primer: 5’-GGGACTTCTCCTACCAAT-3’, reverse primer: 5’-TATATGGTGGATCCTGCAGGGTAG-3’; MUC5B forward primer: 5’-CACATCCACCCTTCCAAC-3’, reverse primer: 5’-GGCTCATTGTCGTCTCTG-3’; ALOX15 forward primer: 5’-GGGCAAGGAGACAGAACTCAA-3’, reverse primer: 5’-CAGCGGTAACAAGGGAACCT-3’). Gene expression was normalized against β-actin and quantified by standard *2^−ΔΔCt^* method.

### Statistical analysis

Prism 8 (GraphPad, San Diego, United States) was used for data presentation and statistical analysis. Data are given as means ± SEM if not stated otherwise. Data were tested for normal distribution using a Shapiro-Wilk test followed by an unpaired two-tailed Student’s *t-test* (normal distribution) or a Welch’s *t-test* (data not normally distributed). Differences between more than two groups were assessed by two-way ANOVA with Tukey’s multiple comparison test or by non-parametric Kruskal-Wallis test. Statistical significance was considered at p < 0.05 (* p = 0.05, ** p = 0.01, *** p = 0.001, and **** p = 0.0001). Experiments were performed in 3-7 independent biological replicates in 2 (qPCR) or 3 (TEER, height) technical replicates (N=3-7, n=2-3). For automated analysis of airway mucus, the entire specimens were scanned and data is given as mean ± SEM (N=12-50 images) per sample.

## Results

### BMGLS does not change the metabolic activity nor alter Calu-3 morphology

We applied BMGLS to the human lung adenocarcinoma cell line Calu-3 over the course of 3 weeks starting with the air-lift of the ALI culture. BMGLS treatment did not affect the morphology of the respiratory Calu-3 epithelium (Figure 1A). The epithelial height remained consistent even at concentrations of 0.6% BMGLS ranging from 61 ± 16 μm to 52 ± 8 μm in the control group (Figure 1B). Furthermore, concentrations of 0.07%, 0.15%, 0.3%, 0.6% 1.25%, 2.5% and 5% (v/v) elicited no cytotoxic effects when compared to 5% HCS (v/v) or the untreated control in Calu-3 cells (Figure 1C). On the one hand, Ki67 positive cells decreased in BMGLS treated samples as opposed to the untreated control. On the other however, no difference was observed in proliferative and apoptotic events compared to the 0.6% HCS treated control as indicated by Ki67 and cPARP staining, respectively, suggesting that decreased Ki67 staining arises from HCS rather than bioactive compounds in BMGLS (Figure 1D).

**Figure 1.**
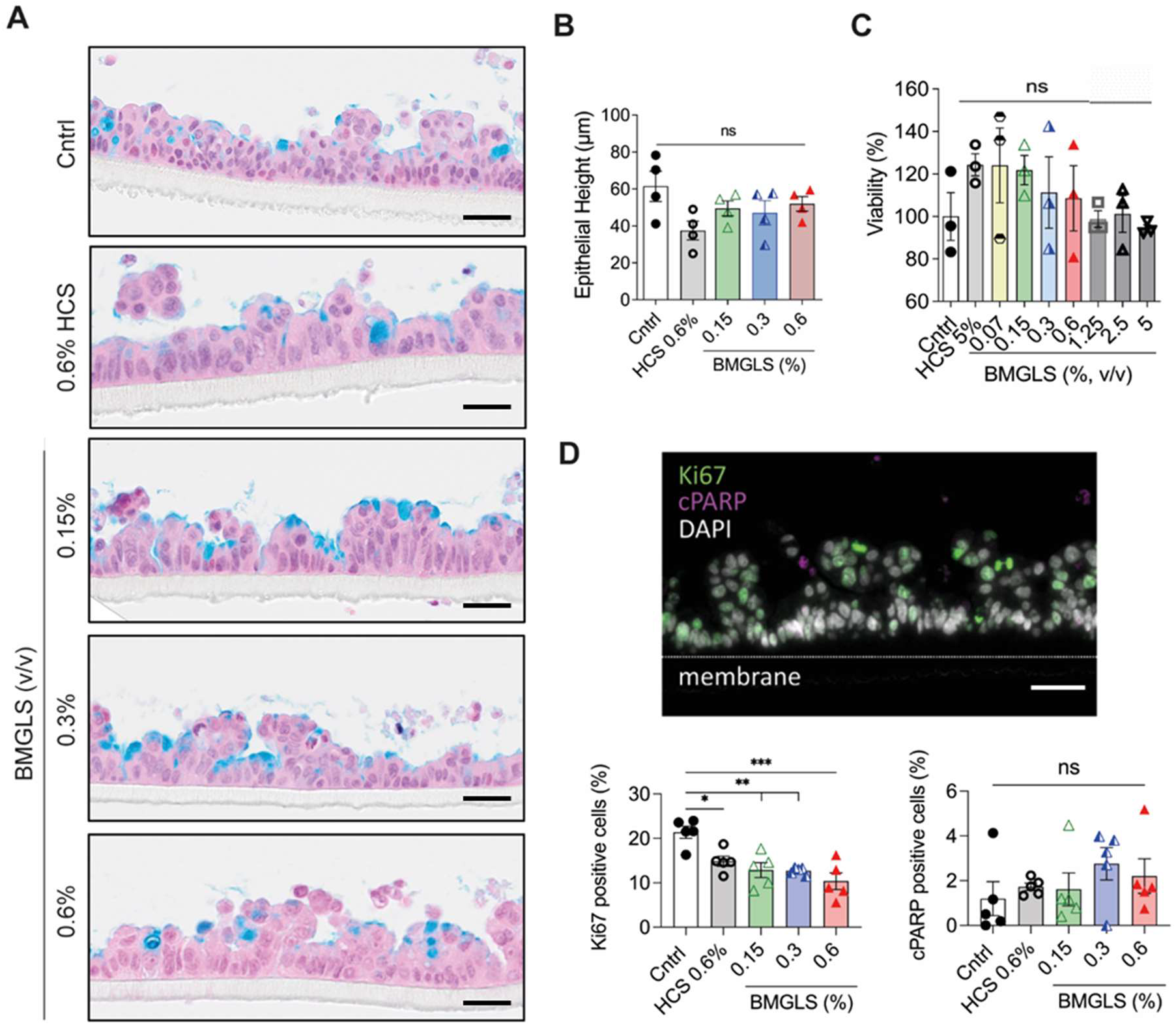
Analysis of BMGLS treated Calu-3 ALI cultures. Calu-3 ALI cultures were treated with different concentrations of BMGLS over the course of 3 weeks. (A) Characterization of cell morphology using AlB staining (acidic mucopolysaccharides in blue; nuclei in purple; cytoplasm in pink). Representative images show no morphological differences between untreated and treated samples. Scale bars 50 μm. (B, C) No significant changes in epithelial height (B) and viability (C) were detected. Values are expressed as mean ± SEM. ns = not significant (Dunnett’s Multiple Comparison). (D) Representative IF image of proliferating cells (Ki67, green), apoptotic cells (cPARP, magenta) and nuclei (DAPI, white). Ki67 positive cells decreased in BMGLS treated samples compared to the control. No difference in the percentage of cPARP positive cells was detected after 3 weeks of BMGLS application. Differences were evaluated by One-way ANOVA with Tukey’s multiple comparison test. Statistical significance was considered at p < 0.05 (*), p < 0.01 (**) and p < 0.0001 (****). Scale bars, 50 μm.

### BMGLS increases metabolic activity in NHBE cells

We assessed a potential cytotoxic effect in primary NHBE cells when exposed to BMGLS by applying the formulation at concentrations of 0.07%, 0.15%, 0.3%, 0.6% 1.25%, 2.5% and 5% (v/v supplied in basal media) for 24h with 5% HCS (v/v supplied in basal media) used as solvent control using an MTT assay (Figure 2). While no effect on viability was measured for HCS at 5%, NHBE cells appeared more viable above BMGLS concentrations of 1.25 % (137 % ± 30 %, p < 0.05) to 5 % (141 % ± 21 %, p < 0.05). However, since the MTT assay measures viability by assessing cellular levels of NAD(P)H, increases might also indicate enhanced metabolic activity. In order to focus on mucosal properties without disturbing any further cellular activities, we chose concentrations of 0.15 % to 0.6 % for further experiments.

**Figure 2.**
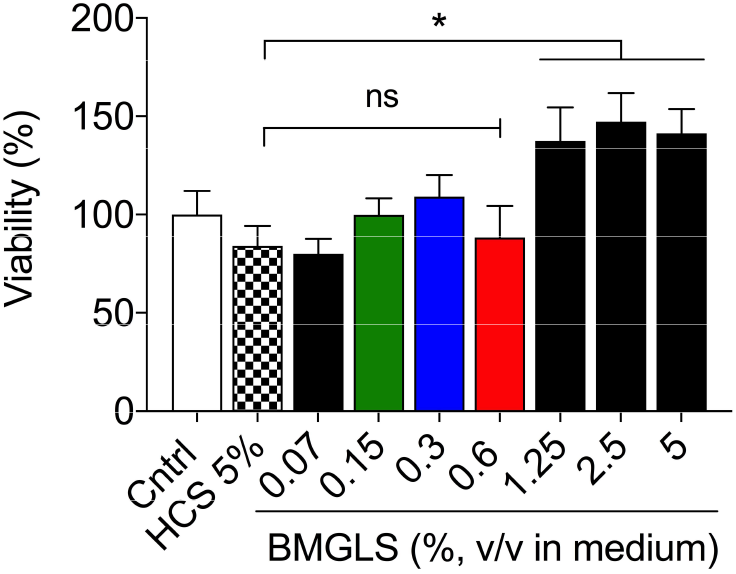
Cell viability measurements to assess cytotoxic effects of BMGLS on primary NHBE cells. NHBE cells were stimulated with concentrations of 0.07 %, 0.15 %, 0.3 %, 0.6 % 1.25 %, 2.5 % and 5 % (v/v) including 5 % HCS as solvent control for 24 h followed by cytotoxicity assessment using an MTT assay. Data was normalized to the unstimulated control. Differences between groups were assessed using One-way ANOVA with Dunnet’s multiple comparisons test. Significance was considered at p < 0.05 (*).

### BMGLS alters NHBE morphology

The isolated primary NHBE cells differentiated into a pseudostratified columnar epithelium within 3 weeks of ALI culture with the presence of apical cilia and mucus producing goblet cells (data not shown). We applied BMGLS to primary NHBE over the course of 3 weeks starting with air-lift of the ALI culture. BMGLS altered the epithelial morphology in a dose-dependent manner by incorporating, herein termed, intra-epithelial structures (IES) that severely altered the pseudostratified morphology still observed in the Cntrl and solvent control (HCS) samples. These IES were filled with mucus or withhold mucus from secretion (representative images shown in Figure 3A). TEER values significantly declined from 461 ± 51 Ω/cm2 (HCS 0.6 %) to 255 ± 96 Ω/cm2 (0.3 % BMGLS) after 7 days and from 604 ± 62 Ω/cm2 (HCS 0.6 %) to 395 ± 46 Ω/cm2 (0.6% BMGLS) after 10 days yet approached Cntrl levels after 21 days of culture (Figure 3B). Although this may indicate a delayed tight junction formation and overall interference of BMGLS with the integrity of cellular barriers, it may also stem from the commencing presence of the mucus-containing IES as BMGLS may force the restructuring and thereby either directly delays tight junction formation or simply prolongs permeability through diffusion within the newly formed liquid-filled IES.

**Figure 3.**
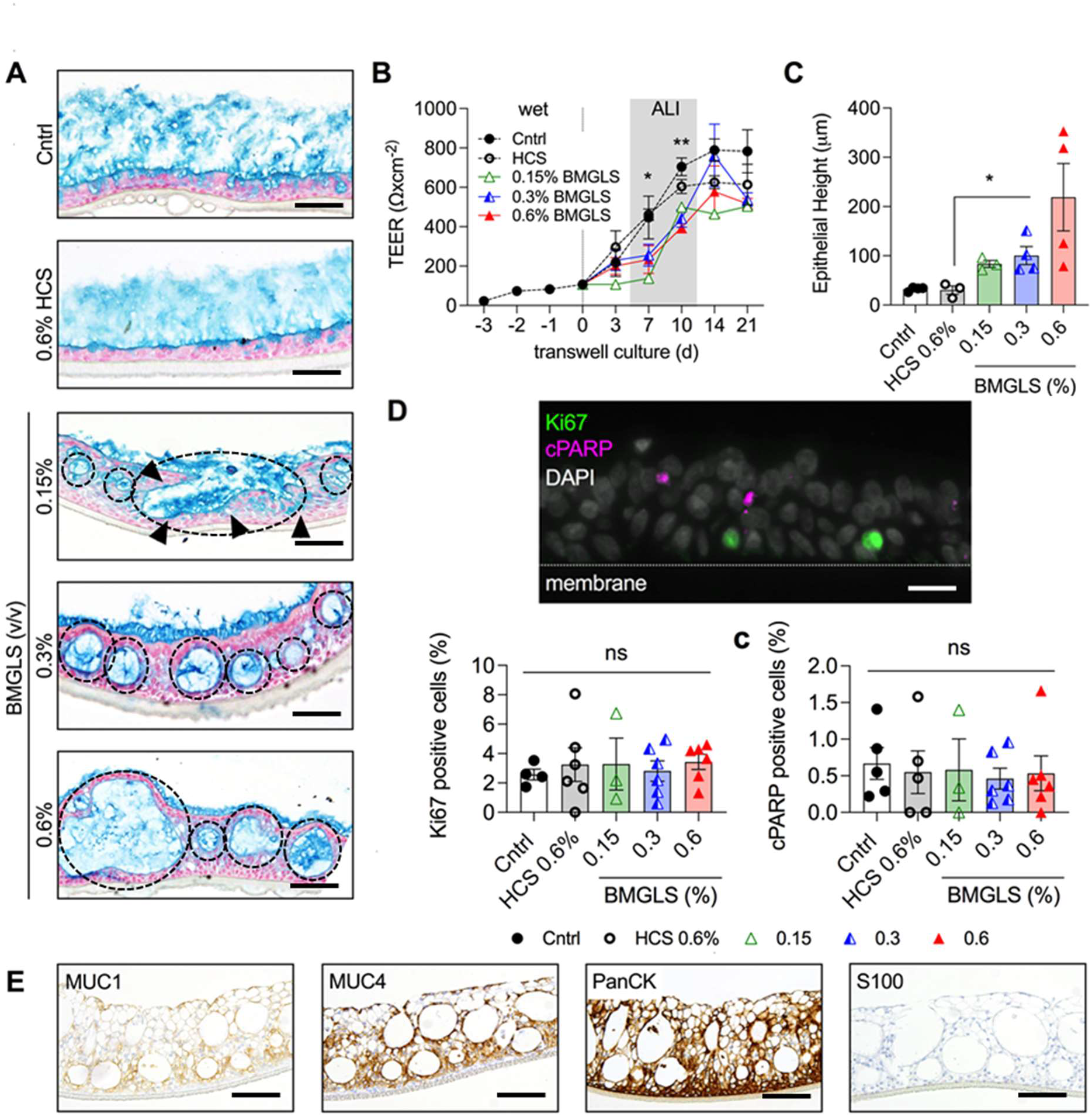
Alterations in NHBE ALI cultures induced by BMGLS. NHBE ALI cultures were supplied with BMGLS over the course of 3 weeks. (A) Increasing concentrations of BMGLS induced the formation of intra-epithelial structures (IES; dotted lines). IES, filled with acidic mucopolysaccharides, identified by using AlB staining (acidic mucopolysaccharides in blue; nuclei in purple; cytoplasm in pink). Black arrows indicate the opening of IES to the surface. Representative images are shown. Scale bars, 200 μm. Analysis was done by TEER (B) as well as histologically (C). Data between groups was compared using (B) One-way-ANOVA with Tukey’s or (C) Dunnet’s multiple comparison test. Significance was considered at p < 0.05 (*), p < 0.01 (**) and p < 0.0001 (****). (D) Representative IF image of proliferating cells (Ki67, green), apoptotic cells (cPARP, magenta) and nuclei (DAPI, white). No difference in the percentage of Ki67 positive or cPARP positive cells was detected after 3 weeks of BMGLS application. Differences were evaluated by One-way ANOVA with Tukey’s multiple comparison test. Statistical significance was considered at p < 0.05 (*). (E) Staining of NHBE ALI cultures following 0.6 % BMGLS application with MUC1, MUC4, and PanCK illustrates epithelial origin and differentiation into a mucus producing REp. S100 stained negative disregarding degenerative processes in NHBE ALI upon BMGLS application. Scale bars, 200 μm.

The morphologic restructuring increased the epithelial height from 32 ± 4 μm in the control group to 219 ± 137 μm (p < 0.05 for 06% BMGLS, Figure 3C). The IES-internal area accounted to 306 ± 219 μm2 in the Cntrl, representing goblet cells, and significantly increased to 2,806 ± 4,646 μm2 at 0.6 % BMGLS treatment (data not shown).

We found no difference between BMGLS treated and non-treated samples in Ki67 or cPARP positive cells overall neglecting influence on proliferation or apoptosis (Figure 3D). Of note, Ki67 positive cells were found in the basal layer of cells supporting the stem cell-like function of basal cells also in the ALI cell culture model.

The observed changes in morphology prompted the need to validate the composition and characteristics of the bronchial epithelial cells. The NHBE ALI culture stained positive for MUC1, MUC4, and PanCK confirming the cellular origin and composition of the bronchial epithelial cells. Furthermore, the typical tumor marker S100 tested negative suggesting to disregard cellular degeneration (Figure 3E).

### SEM and TEM analysis of altered NHBE morphology

The severe changes in morphology were further ultra-structurally analyzed using SEM. The NHBE ALI cultures under Cntrl and HCS conditions reveal the previously observed pseudostratified conformation with consistent lining of apical cilia and embedded secretory cells with microvilli and densely packed mucus (Figure 4A, B). Secreted mucus bundles nicely represent the reported supra-molecular topology of these thread-like highly entangled MUC5B polymers (Figure 4A.ii) coated with MUC5AC (Figure 4A.iii). HCS application to NHBE ALI cultures did not affect cellular morphology, cilia formation or mucus production (Figure 4B). IES were also consistently discovered in SEM samples. Astoundingly, we spotted several IES that seemed to open towards the apical surface which would allow disgorging the secreted mucus from the IES under the caveat that functional cilia would indeed line the IES (Figure 4C), as previously implied by IF. In contrast to the Cntrl, the mucus constrained within these IES displayed loose packing (Figure 4C, close up view left corner). IES in the 0.6 % BMGLS sample were enlarged and support tubular-like conformation with secreted mucus emerging from the breakage that resulted from SEM processing (Figure 4D.i) with otherwise untampered mucin network organization post-secretion across all samples (Figure 4C.ii, D.ii). A potential orifice of an IES is shown in Figure 4D.iii.

**Figure 4.**
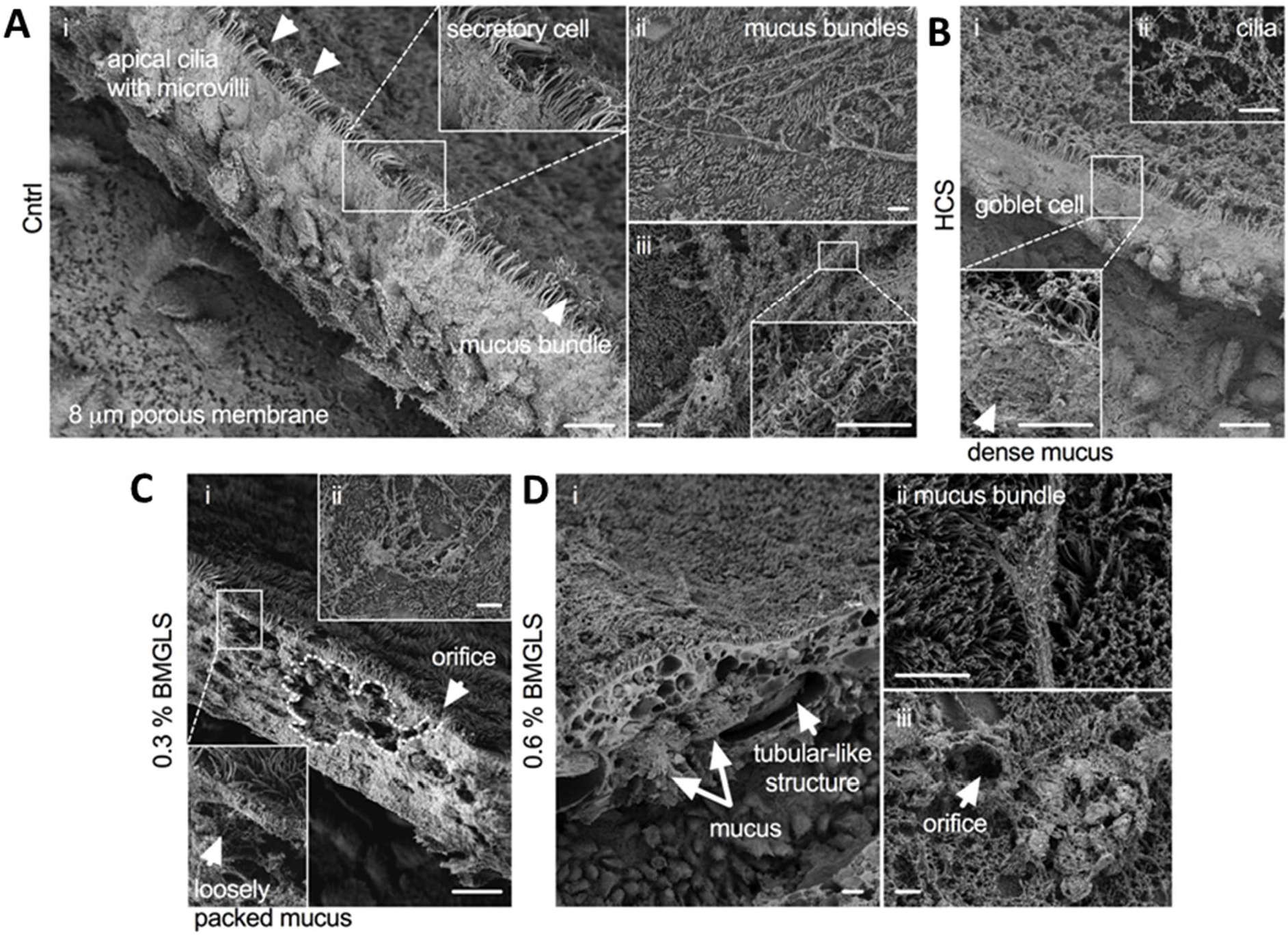
SEM reveals expanded surface area and loosely packed mucus in IES. NHBE ALI cultures after 3 weeks of BMGLS application and processing for SEM. (A) Cntrl sample with (i) pseudostratified morphology and apical cilia. (ii) Mucin bundles and (iii) close-up view of MUC5AC covering MUC5B entangled polymers. (B) HCS sample with densely packed mucus (inset) and (ii) apical cilia. (C.i) IES with apical orifice and loosely packed mucus (inset) at 0.3 % BMGLS application and (ii) secreted mucin bundles. (D.i) 0.6 % BMGLS induces enlarged IES comprising mucus within tubular-like structures. (ii) Apical mucin bundles and (iii) orifice from apical point of view. Scale bars, all 10 μm.

Moving to TEM analysis, goblet cells in NHBE ALI cultures exposed to HCS (0.6 %, v/v) were of normal appearance with electron dense granules and apical microvilli (Figure 5A). However, samples exposed to BMGLS consistently showed the observed IES (Figure 5B, #) with longitudinally cut mucin bundles and intra-epithelial cilia (Figure 5B, inset; c, cilia), whose overall appearance and content was highly distinguished compared to mucosal granules encased within a goblet cell (*). These mucosal granules appeared tumid but with reticularly distributed electron density (Figure 5D2).

**Figure 5.**
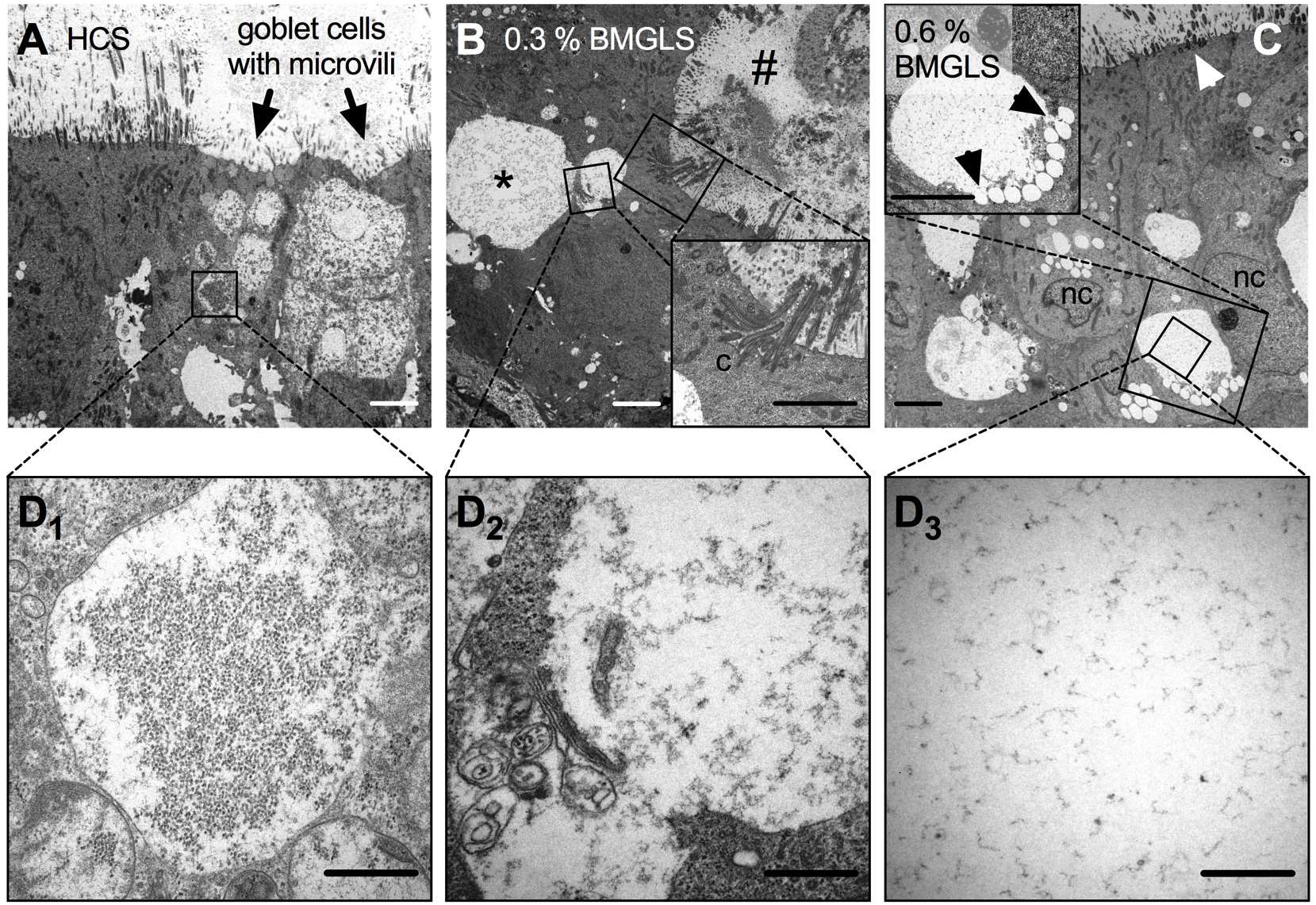
TEM reveals dispensed reticular appearance of IES content. NHBE ALI cultures were treated with BMGLS at different concentrations and processed for TEM. (A) Samples treated with HCS (0.6 %, v/v) show goblet cells with apical microvilli and mucosal granules of reticular appearance, and amorphous electron-dense content (D1). (B) 0.3 % BMGLS (v/v) induces IES formation (#) with internal cilia (c, close-up view) and goblet cells (*) with decreased electron density (D2). (C) Merging of electron- lucent vesicles with mucus granules in 0.6 % BMGLS (v/v) treated samples (close-up view) substantially decreases electron density in the granular content (D3). Scale bars, 5 μm in A-C, and 1 μm in D1-D3.

We further noticed several electron-lucent vesicles merging with a mucosal granule, supposedly discharging its content (Figure 5C; arrows indicating vesicle fusion; nc, nucleus). Overall, the density of the amorphous network in mucosal granules of HCS samples seems to decrease with increasing concentrations of BMGLS supporting findings on its re-hydrational effects (Figure 5D1-D3).

### BMGLS modifies mucus density and composition

To further study mucous properties under the influence of BMGLS, we scanned large areas of the samples and found that mucus for most of the part detached from the PCL in 0.3% and 0.6% BMGLS samples, either completely or it accumulated at distant locations (Figure 6A; arrows indicating PCL and detached mucus). Computational analysis revealed that the mucosal area in these samples decreased significantly when normalized to the membrane length (Figure 6B). Considering AlB intensities, we observed an unexpected increase at a concentration of 0.15% suggesting higher rather than lower mucus density (Figure 6C). However, the AlB intensities showed a trend (p < 0.08) towards reduced levels with increasing concentrations of BMGLS, potentially indicating mucus swelling.

**Figure 6.**
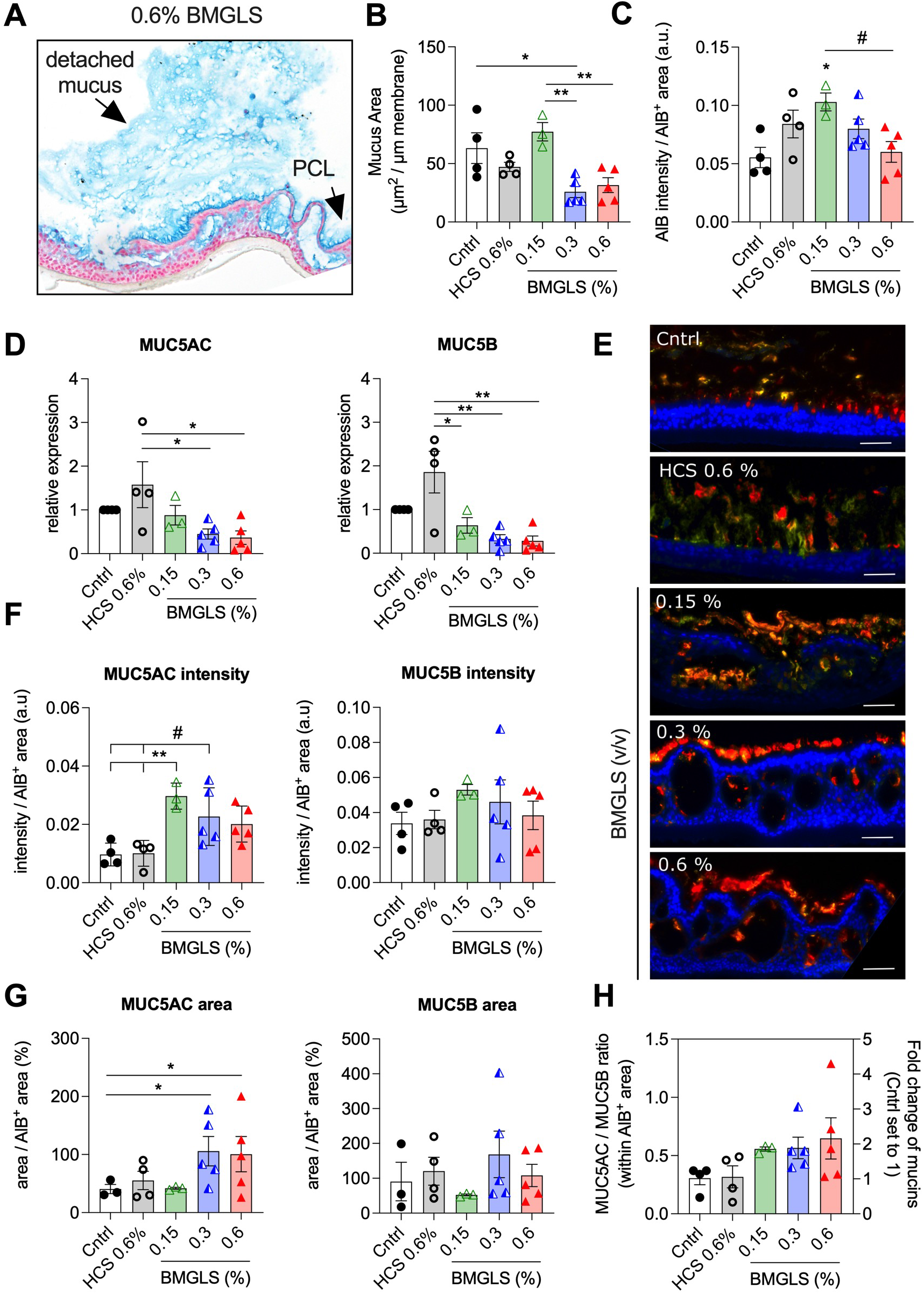
BMGLS facilitates detachment of the mucus layer from the PCL. BMGLS treated NHBE ALI cultures were analyzed for (A, B) the mucus area (given in μm2 per μm membrane length), (C) the AlB intensity (a.u.) per AlB+ area, and (D) the relative expression of MUC5AC and MUC5B mRNA. The mucus layer detached from the PCL (A, arrows) and accumulated at different locations across the sample. Differences between groups were computed by One-way ANOVA with Tukey’s multiple comparison test. Significance was considered at p < 0.05 (*) and p < 0.01 (**) with (#) indicating a trend at p < 0.08. (E) Representative images of MUC5AC and MUC5B in AB-IF double staining. NHBE ALI samples were treated with BMGLS at different concentrations over 3 weeks and histologically processed for AlB-IF based computational analysis targeting MUC5AC (red) and MUC5B (yellow). Nuclei are stained using DAPI (blue). Scale bars, 20 μm. (F-H) BMGLS attenuates MUC5AC in airway mucus.

Hence, we speculate that the swelling, probably mediated by rehydration, causes the mucus to detach from the PCL, leading either to loss of the mucus explaining the reduced mucosal area, or mucus accumulation at different positions across the sample. Overall, this might indicate that under applied cell culture conditions, BMGLS stimulates mucus production at lower concentrations while fostering rehydration when supplied at higher dosages.

Next, we evaluated expression levels of MUC5AC and MUC5B including their ratio using RT-qPCR and our recently developed image-based analysis approach to assess protein levels (Groiss et al., 2022). BMGLS application significantly reduced mRNA expression of both MUC5AC and MUC5B (Figure 6D). Representative color images of all merged channels are shown in Figure 6E. The reduction in gene expression did not correlate to protein levels as the MUC5AC intensity increased at 0.15 % BMGLS (v/v) while MUC5B protein levels remained stable across all samples (Figure 6F) leading to a 2-fold increase in the MUC5AC/MUC5B ratio (Figure 6H). We further used the pipeline to calculate the MUC5AC^+^ and MUC5B^+^ areas within the total mucosal area. The MUC5AC^+^ area enlarged significantly in samples stimulated with 0.3 % and 0.6 % BMGLS supporting previous results on mucus swelling (Figure 6G). This was not observed for the MUC5B^+^ area maybe because its entangled polymeric conformation hinders dissolution in applied conditions (Figure 6G).

### BMGLS targets ALOX15

Lastly, we tested interference of BMGLS with the major mucus-stimulating enzyme ALOX15. BMGLS significantly reduced ALOX15 gene expression at all tested concentrations when compared to the solvent control (HCS 0.6 %, v/v; Figure 7A). Upon evaluating whether this finding translate to protein levels, we found diminished levels of ALOX15 in BMGLS treated samples (Figure 7B).

**Figure 7.**
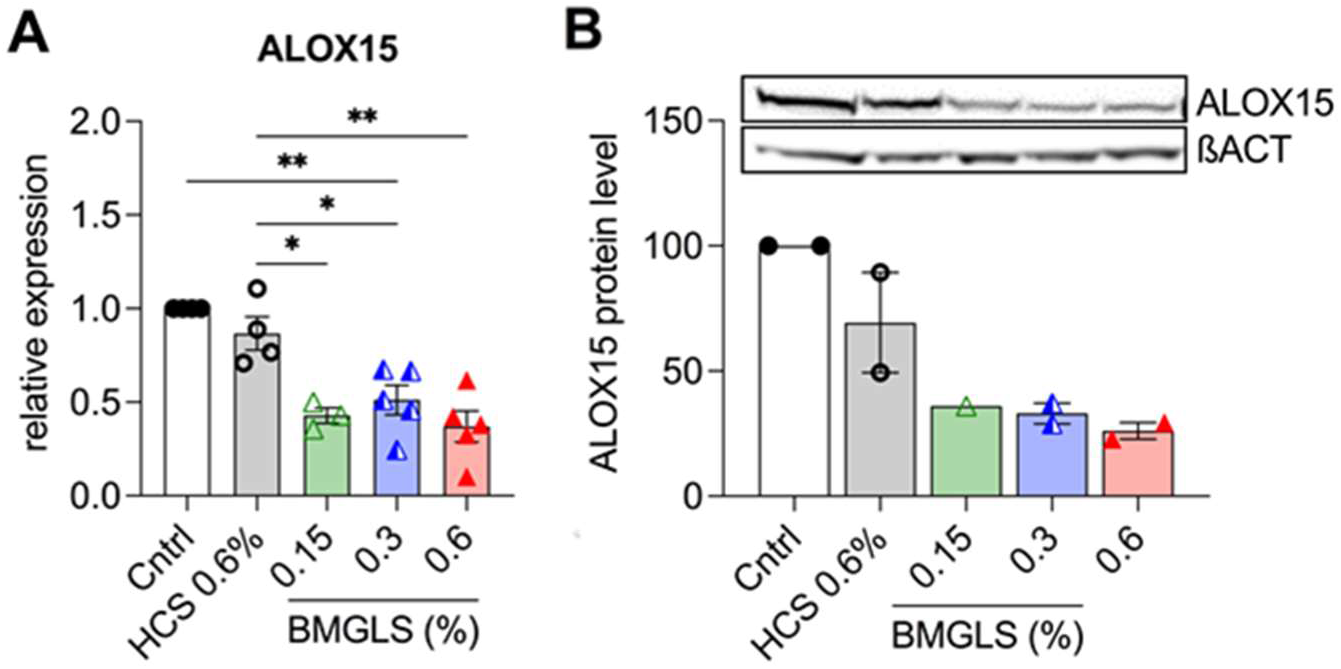
BMGLS reduces expression of the mucus-stimulating enzyme ALOX15. Gene expression (A) and protein expression (B) of ALOX15 in NHBE ALI cultures treated with BMGLS for 3 weeks. Differences between groups were computed by One-way ANOVA with Tukey’s multiple comparison test. Significance was considered at p < 0.05 (*) and p < 0.01 (**).

## Discussion

Medicinal herbs are widely used to treat diverse respiratory discomforts in Southeast Asian and Western Pacific areas (Yang et al., 2018) and proved beneficial in the treatment and containment of COVID-19 (Luo et al., 2020). Especially members of the *Fritillariae* family that also constitute the main ingredient of BMGLS become increasingly attractive to western medicine in order to better understand their pharmacodynamics, pharmacokinetics and toxicology, but most importantly, their potential therapeutic effects for use within CRD treatment (Chen et al., 2020; Li et al., 2019).

In our studies, we observed severe remodeling of the reconstituted REp under the influence of BMGLS. The formation of IES induced an increase in surface area primarily within the IES marked by the presence of cilia and encasing of mucus and was dose-dependent. IES appeared to fuse and form large tubular-like structures with orifices towards luminal areas potentially facilitating mucus release from IES. Although the detailed mechanism of airway remodeling remains unclear, this process is associated with severe or persistent inflammation in COPD (Wang et al., 2018). Since the REp cultures in our experiments were established from primary NHBE cells, they do not contain the necessary inflammatory cells to induce such airway remodeling. Nevertheless, the apparent changes are very intriguing and are probably attributed to GCM and GCH as well as potential dedifferentiation of basal or club cells present within the reconstituted REp.

The observed decrease in TEER values in BMGLS-treated samples at day 7 and 10 of differentiation potentially indicates a slower formation of tight junctions (Kojima et al., 2013; Srinivasan et al., 2015). However, since no differences in TEER were observed after the NHBE ALI culture was fully established, we speculate that BMGLS postpones barrier closure by stimulating a diverse set of cellular effects rather than directly decreasing tight junction formation itself. The reduced TEER may also be caused by substantial cell-free areas within these IES, which might facilitate the flow of water, solutes and electrolytes (Flynn et al., 2009). Further, staining of airway epithelial marker such as PanCK, MUC1, MUC4 and S100 confirmed cellular origin and neglected cellular degeneration. On the other hand, we ascertained the internal area of IES to be indeed surface area with cilia formation and expression of mucins. Furthermore, the reduced occurrence of apical cilia found in SEM images apparently suggests increased re-differentiation of ciliated to mucus producing goblet or club cells implying a potential stimulating effect of BMGLS toward GCM (Boucherat et al., 2013; Bustamante-Marin & Ostrowski, 2017). Whether similar effects also co-occur in the fully established REp when treated with BMGLS within the same three-week period remains to be investigated. Hence, BMGLS appears to remodel the establishing REp without executing overall degeneration potentially by altering cellular differentiation which causes the observed changes in morphology.

Hyperproduction or hyperdensity of airway mucus impairs effective MCC and thereby constitutes a major contributor to all known airway diseases. Pathologic conditions caused by excessive exposure to environmental insults not only increase the number of solids present in airway mucus, but also change the compositions and ratio of the MUC5AC and MUC5B that is essential for airway homeostasis (Fahy et al., 1997; Rose & Voynow, 2006; Roy et al., 2014). Compounds of *Fritillariae* species, precisely imperialine, chuanbeinone, peimine and peiminine, have been shown to exhibit expectorant properties in mice measured by tracheal phenol red output (Wang et al., 2011). Similarly, imperialine-β-N-oxide, isoverticine and isoverticine-β-N-oxide increased tracheal phenol red output in mice using a model of ammonia-induced cough (Wang et al., 2012). Hence, the expectorant effects observed in *Fritillariae* species may attribute to the plants potential to increase trachea-bronchial mucus secretion.

Based on these findings, we hypothesized that the reported increase in secretion is due to changes in the hydration level of airway mucus, either by reducing the protein content or by directly increasing hydration rates. Both of these processes would facilitate detachment of the mucous layer from the PCL and allow for mucosal clearance.

While BMGLS reduced the mRNA expression levels of both MUC5AC and MUC5B as well as the mucus area in our study, it seemed to increase the AlB intensity at 0.15 % (v/v), which returned to baseline only at higher dosages. Given that AlB intensity correlates with the presence of mucopolysaccharides hence the amount of mucins, these findings would actually indicate an increase in mucins and hence mucus density that seems antagonized by elevated hydration at higher dosages. Intriguingly and in contrast to the observed mRNA expression levels, BMGLS seemed to induce MUC5AC protein expression increasing the MUC5AC/MUC5B ratio by 2-fold. However, the area occupied by MUC5AC enlarged, overall corroborating findings on mucus swelling that might underlie the dispersion of the surplus MUC5AC.

The notion that BMGLS reduces the mucous area most likely by facilitating mucus detachment from the PCL was supported by the discovery of substantial mucus accumulations at the margins of the samples (Figure 6A, B). These were not visible in the Cntrl or HCS as the mucus in these samples was distributed evenly throughout the membrane (Figure 3A), strongly advocating that mucus indeed detaches from the PCL under the influence of BMGLS. Particularly the complete detachment of mucus would explain the reduced mucus area in our automated analysis. The similar levels of AlB intensity at 0.6 % BMGLS and the Cntrl might therefore stem from the remaining PCL that is less subject to increased rates of hydration, consequently staining at the same AlB intensity as the Cntrl (Bustamante-Marin & Ostrowski, 2017).

Furthermore, we observed several electron lucent vesicles merging with a large mucosal granule in TEM images (Figure 5C). The merging of secretory vesicles for combined secretion is known as “compound exocytosis” and is well described in zymogen granules in pancreatic acinar cells (Hansen et al., 1999; Pickett & Edwardson, 2006; Takahashi & Kasai, 2007) and intestinal goblet cells (Birchenough et al., 2015; Grootjans et al., 2013; Liu et al., 2015). During this event, multiple vesicles fuse and rapidly disgorge their content into the luminal area (Birchenough et al., 2015). Sequential compound exocytosis of secretory granules was also reported in the nasal cavity (Oshima et al., 2005). However, mechanistic studies are still missing as are reports describing this phenomenon in goblet cells of the proximal airways (Adler et al., 2013).

It is uncertain whether the observed events in our study indeed attribute to compound exocytosis. Nevertheless, disgorging the vesicular content into the larger granules seemed to disperse the amorphous mucous network within (Figure 5D_1_-D_3_). Hence, BMGLS might function as secretagogue by inducing compound exocytosis-like processes in the REp in order to increase hydration of mucus for facilitated secretion and release from the PCL.

Overall, BMGLS seemed to enhance hydration rates of airway mucus via so far unidentified processes that facilitate detaching of the mucous layer from the PCL, which is a prerequisite for efficient expectoration. Exocytosis-like events might explain the enlarging of intracellular granules, however, they are unable to account for the formation of mucus-containing IES. Ultimately, more investigations are needed to identify a potential mechanistic link between morphologic changes and mucus production and secretion.

An increasing line of evidence suggests the involvement of ALOX15, including its metabolite 15-hydroxyeicosatetraenoic acid (15-HETE), in GCM and mucus hyperproduction in pathologic conditions such as asthma (Zhao et al., 2009). ALOX15 has recently been suggested as part of a predictive gene signature in exacerbations of COPD patients to personalize treatment decisions (Ditz et al., 2021). On the other hand, enhanced levels of ALOX15 expression have been reported to accompany the differentiation of NHBE cells at ALI conditions despite the lack of external stimulation, suggesting that baseline levels of ALOX15 are essential for physiologic production of airway mucus (Jayawickreme et al., 1999).

ALOX15 mRNA expression levels decreased significantly at all tested concentrations of BMGLS in comparison to the HCS control in our study. This data was supported by reduced ALOX15 protein expression suggesting that BMGLS might ease mucus production by lowering levels of the mucus-stimulating enzyme ALOX15. The ALOX15 pathway was also shown to be inhibited by baicalein and coptisine, the bioactive ingredients of another TCM formulation with anti-inflammatory properties known as Huang Lian Jie Du Tang (HLJDT) (Zeng et al., 2011). Therefore, most likely there is a wide range of medicinal plants used in TCM that hold the potential for treatment of CRDs.

A major limitation of our study is the use of hydrophilic concentrates directly within our cell culture experiments. BMGLS is usually applied orally, which means that the bioactive compounds need to be stable enough for the passage through the gastrointestinal tract or that its ingredients may actually be converted to the ultimate bioactive product by the intestinal microflora (Gao et al., 2019; Houriet et al., 2021; van Dooren et al., 2018).

Therefore, future studies should aim at investigating the *in vivo* effects of the digested formulation in order to correlate bioactivities observed *in vitro* with the efficacy and potential toxicology *in vivo*.

## Conclusion

Initial insights suggest that BMGLS may constitute an effective secretagogue by stimulating mucus production in combination with increasing mucus hydration, which overall promotes detachment of the mucous layer from the PCL for facilitated expectoration. In addition, compound exocytosis-like processes in goblet cells and the regulation of mucus stimulating pathways such as ALOX15 seem to be part of the potential underlying mechanism that mediate the observed findings. BMGLS has been used in the treatment of CRDs in TCM over two millennia without reports on adverse effects on the human health. Nevertheless, further investigations on toxicology and dosimetry are called for to clarify the unexpected morphological changes in the reconstituted REp before proceeding to *in vivo* studies.

## Supplemental Material and Methods

### Characterization of NHBE cells after isolation

We confirmed the viability/ quality of isolated NHBE cells immediately after isolation by trypan blue staining and Casy Cell Counter measurements. To check identity of cells we used IF to verify individual cell types in the re-established pseudostratified epithelium. αTubulin (αTub) was used to target apical cilia while MUC5AC and uteroglobin (CCSP) mark the presence of goblet and Club cells, respectively. Cilia formation was present consistently in NHBE ALI cultures after 3 weeks of differentiation similarly to MUC5AC (representative images shown in Supplemental Figure 1 a, b). Interestingly, CCSP was detected only scarcely in normal NHBE ALI cultures (Supplemental Figure 1 a, b; white arrows).

**Supplemental Figure 1.**
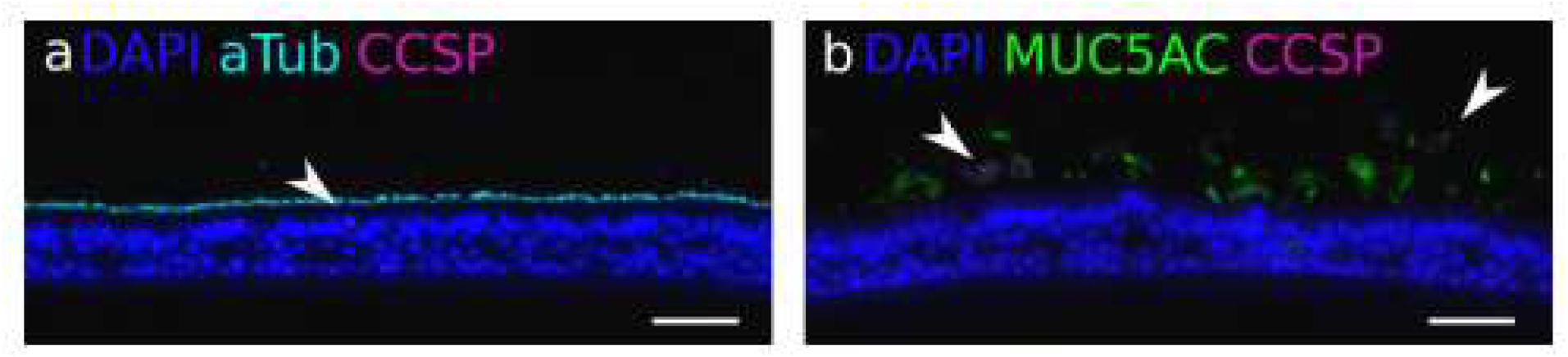
Expression of αTub, CCSP and MUC5AC in ALI cultures. Representative images of (a) αTub that was expressed comprehensively across the sample and (b) MUC5AC and CCSP, marking the presence of goblet and club cells, respectively. CCSP was present only in sparse amounts (a, b; arrowheads). Scale bars, 20 μm.

